# The clade-specific target recognition mechanisms of plant RISCs

**DOI:** 10.1101/2024.01.03.574122

**Authors:** Hiro-oki Iwakawa

## Abstract

Eukaryotic Argonaut proteins (AGOs) assemble RNA-induced silencing complexes (RISCs) with guide RNAs that allow binding to complementary RNA sequences and subsequent silencing of target genes. The model plant *Arabidopsis thaliana* encodes 10 different AGOs, categorized into three distinct clades based on amino acid sequence similarity. While clade 1 and 2 RISCs are known for their roles in post-transcriptional gene silencing, and clade 3 RISCs are associated with transcriptional gene silencing in the nucleus, the specific mechanisms of how RISCs from each clade recognize their targets remain unclear. In this study, I conducted quantitative binding analyses between RISCs and target nucleic acids with mismatches at various positions, unveiling distinct target binding characteristics unique to each clade. Clade 1 and 2 RISCs require base pairing not only in the seed region but also in the 3′ supplementary region for stable target RNA binding, with clade 1 exhibiting a higher stringency. Conversely, clade 3 RISCs tolerate dinucleotide mismatches beyond the seed region. Strikingly, they bind to DNA targets with an affinity equal to or surpassing that of RNA, like prokaryotic AGO complexes. These insights challenge existing views on plant RNA silencing and open avenues for exploring new functions of eukaryotic AGOs.

## Introduction

Argonaute family proteins bind to target nucleic acids through small RNA or DNA guides and are involved in the exclusion of non-self nucleic acids and accurate gene expression regulation. AGOs are present in all domains of life, highlighting their fundamental role (1–3). Eukaryotic AGOs form RNA-induced silencing complexes (RISCs) with non-coding RNAs of ∼20–30 nucleotides (nts) as guides. Through these guides, they interact specifically with target RNAs, regulating gene expression at the transcriptional or post-transcriptional level (Figure 1A) (4). In contrast, prokaryotic AGO proteins can use not only small RNAs but also small DNAs as guides. They primarily bind to partially melted single-stranded DNA (ssDNA) regions (Figure 1A) and are known to play roles in cleavage of foreign DNA such as plasmids, transposons or phage DNAs and in unlinking chromosome catenanes in the absence of gyrase (2, 5). Generally, AGOs consist of six domains: N, Linker 1 (L1), PAZ, L2, MID, and PIWI (2, 6). Structurally, they adopt a bilobed configuration composed of MID-PIWI and N-PAZ domains. The 5′ end of the guide strand binds to a pocket formed by the MID domain, while the 3′ end binds to the PAZ domain (7–9). The PIWI domain exhibits an RNaseH-like fold and, depending on the type of AGO, possesses activity to cleave the target RNA or DNA that forms base pairs with the guide nucleic acid (10–12).

**Figure 1.**
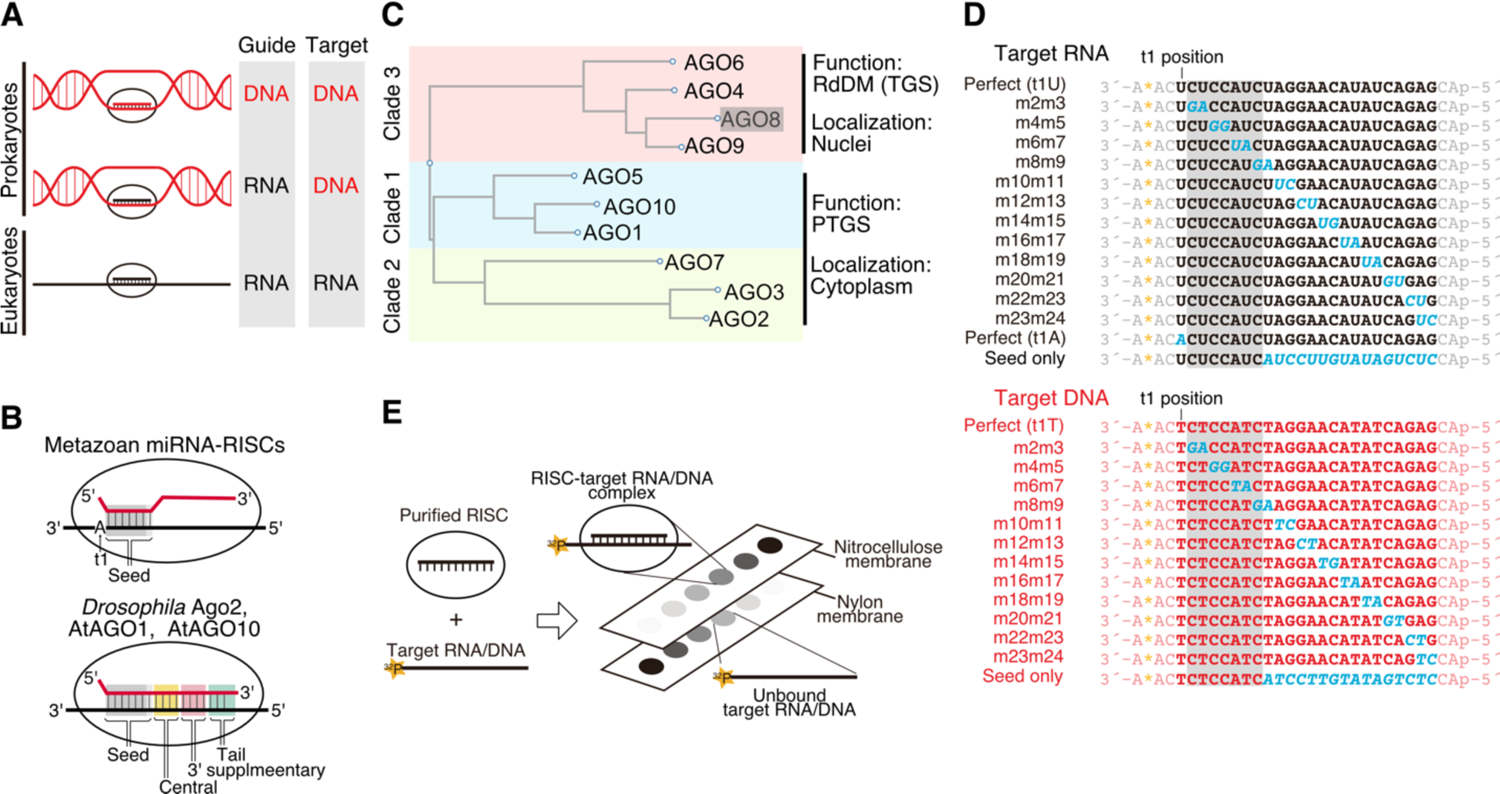
Analysis of target binding in three clades of RISCs in *Arabidopsis thaliana*. A. The main combinations of guide nucleic acids and target nucleic acids in prokaryotic and eukaryotic AGOs. B. Base pairing required for target binding by metazoan miRNA-RISC, and RISCs specialized in target cleavage (e.g., *Drosophila* Ago2-, AtAGO1- and AtAGO10-RISCs). Base pairs corresponding to the seed region, central region, 3′ supplementary region, and tail region are highlighted. C. Phylogenetic tree of Arabidopsis AGOs. AGO1, 5, 10 belong to clade 1, AGO2, 3, 7 belong to clade 2, which mainly induce PTGS in the cytoplasm. Within clade 3, AGO4, 6, 8, 9, of which AGO8 is a pseudogene, the others mainly induce RNA-directed DNA methylation (RdDM) in the nucleus. D. Target RNAs and DNAs used in this study. Italicized letters indicate mutation sites. The letter A* denotes cordycepin (^32^P). The gray shadow indicates the seed region. E. Schematic of the filter binding assay used in this study.

Research on eukaryotic AGOs has revealed that when guide RNA forms RISC with AGO, the guide RNA is divided into four domains: the seed (from the second nucleotide to the eighth nucleotide of the guide RNA: g2–g7 or g2–g8), central (g9–g12), 3′ supplementary (g13–g16), and tail regions (g17–g21) (Figure 1B) (4, 13). The formation of base pairs between the seed region and the target sequence (t2–t7 or t2–t8) is essential for the interaction between RISC and the target (14–17). The interaction between the 3′ supplementary region and the target sequence (t13-t16) have a role in assisting binding at the seed region (17–21). Furthermore, introducing mismatches into the tail and central regions does not significantly affect binding affinity, but mismatches in the central region inhibit target cleavage (13, 19). Metazoan microRNA (miRNA)-RISC can bind to the target using only the seed region (Figure 1B) and suppress target gene expression by recruiting factors that destabilize mRNA through interaction with the scaffold protein GW182, rather than cleaving the target (4, 13, 22). In contrast, Ago2-RISC in *Drosophila* (DmAgo2) and *Arabidopsis* AGO1- and AGO10-RISCs require more extensive base pairing for stable binding and target cleavage (Figure 1B) (13, 23, 24). Additionally, as mentioned earlier, prokaryotic AGO primarily binds to DNA and performs functions such as plasmid cleavage. Thus, the target binding property of RISC reflects its function.

Plants encode multiple AGOs with distinct roles. The model plant, *Arabidopsis thaliana*, encodes ten different AGOs classified into three clades based on amino acid sequence similarity (Figure 1C) (25). AGO4, AGO6, and AGO9, which belong to clade 3 along with AGO8, known to be a pseudogene, incorporate 24-nt small interfering RNAs (siRNAs) in the cytoplasm and subsequently bind to the newly synthesized transcripts of the plant-specific RNA polymerase Pol V in the nucleus (26–30). These clade 3 RISCs recruit the DNA methyltransferase DOMAIN REARRANGED METHYLTRANSFERASE 2 (DRM2), promoting sequence-specific DNA methylation, a mechanism known as RNA-directed DNA methylation (RdDM), which leads to transcriptional gene silencing (TGS) (26, 31). In contrast, AGO1, AGO5, and AGO10, which belong to clade 1, form complexes with 20–22 nt microRNAs (miRNAs) and siRNAs to primarily carry out target RNA cleavage and translation inhibition in the cytoplasm (23, 27, 30, 32–35). Clade 2 AGOs (AGO2, AGO3, and AGO7) also mainly induce post-transcriptional gene silencing (PTGS) in the cytoplasm (30, 36–39). However, there have also been reports indicating that clade 1 and clade 2 AGOs can function in transcription activation, DNA repair, and DNA methylation in the nucleus (40–42).

In this study, I aimed to elucidate the target recognition properties of plant RISC from each clade by quantitatively examining their interactions with targets. I measured the binding affinities between AGO1 (clade 1), AGO2 (clade 2), AGO4 (clade 3), each programmed with small RNAs, and a variety of target RNAs/DNAs with various mutations *in vitro*. The findings revealed distinct target binding properties for RISC in each clade. Clade 1 and clade 2 RISCs required stable base-pairing in the 3′ supplementary region in addition to the seed region for efficient target binding. In contrast, clade 3 RISC tolerated dinucleotide mismatches beyond the seed region. Surprisingly, although clade 1 and clade 2 RISCs prefer RNA over DNA, clade 3 RISCs showed higher binding affinity to DNA than to RNA. These results not only prompt a reconsideration of current models of plant RNA silencing but also unveil previously undiscovered functions of eukaryotic RISCs.

## Methods

### Plasmids

pBYL-3×FLAG-SUMO-catalytic mutant AtAGO2, 4, 6, 9 were created by replacing the catalytic mutant AtAGO1 part of the previously developed pBYL-3×FLAG-SUMO-catalytic mutant AtAGO1 (AtAGO1^D762A^) (23) with sequences coding for the catalytic mutant variants of each respective AGO (AtAGO2^D745A^, AGO4^D660A^, AGO6^D623A^, AGO9^D632A^).

### *In vitro* transcription

The mRNA coding for the catalytic mutant AGOs used in the equilibrium binding assay was transcribed *in vitro* using the AmpliScribe T7 High Yield Transcription Kit (Lucigen), employing the corresponding pBYL-3×FLAG-SUMO-catalytic mutant AGO plasmids linearized with Not I as templates. These mRNAs were subsequently capped using the ScriptCap m^7^G Capping System (Cell Script).

### Lysate preparation

The method for preparing lysate from tobacco BY-2 cells has been detailed previously (43). Briefly, tobacco BY-2 cells on the third day of culture were treated with cellulase and pectolyase to form protoplasts. These were then subjected to Percoll density gradient centrifugation to remove vacuoles. The cells were subsequently lysed using a Dounce homogenizer, and the supernatant was collected after centrifugation at 20, 000×g.

### RISC assembly and purification

The protocol for RISC assembly and purification using a cell-free system has been detailed previously (23). Briefly, 500 µl of BY-2 lysate was mixed with 250 µl of substrate mixture and 50 µl of 1 µg/µl 3×FLAG-SUMO-catalytic mutant AGO mRNA, and incubated for 30 minutes at 25°C. Subsequently, 100 µl of 1.5 µM small RNA duplex was added, and the mixture was incubated for another 1.5 hours at 25°C. The reaction mixture was incubated with anti-FLAG antibody (Sigma), immobilized on Dynabeads protein G (Invitrogen), for 60 minutes at 4°C. Subsequently, the beads were subjected to three washes with lysis buffer (30 mM HEPES (pH 7.4), 100 mM KOAc and 2 mM Mg(OAc)_2_) containing 0.1% Triton X-100. Afterward, the beads were further incubated in equilibration buffer (100 mM KOAc, 18 mM HEPES at pH 7.4, 3 mM Mg(OAc)_2_, 5 mM DTT, 0.01% (v/v) IGEPAL CA-630, and 0.01 mg/ml baker’s yeast tRNA (Sigma)) supplemented with 20% glycerol and 0.05U/μl SUMOstar protease (LifeSensors) for 20 minutes at 4°C to elute the catalytic mutant AtAGO-RISCs.

For the assembly of AGO1-RISC, a 21-nt small RNA duplex with U at the 5′ end of the guide strand was utilized, while for AGO2-RISC assembly, a 21-nt small RNA duplex with A at the 5′ end of the guide strand was employed (Supplementary Figure 5). To ensure the preferred incorporation of the targeted strand as the guide strand, these small RNA duplexes were designed with a mismatch at position 1. Furthermore, to facilitate the ejection of the passenger strand, mismatches were introduced at positions 5 and 10 in the duplexes. For the assembly of clade 3- RISC, a 24-nt small RNA duplex with A at the 5′ end of the guide strand was used (Supplementary Figure 5). This duplex had mismatches at positions 5, 9, and 16 to enhance passenger strand ejection. It should be noted that although this duplex lacks a mismatch at position 1, the guide strand selection is preferentially biased, as one strand’s 5′ terminal base is A while the other strand’s is G, as described by Liu et al., 2022 (44).

### Quantification of RISC using splint ligation method

The concentration of RISC was estimated based on the concentration of the guide strand incorporated into AGO. Quantification of the guide strand was performed using the splinted ligation method (45). After deproteination of the purified RISC with Proteinase K, ethanol precipitation was conducted to extract the guide strand. It was then mixed with a 5′-end-labeled ligation oligo (5′-CGCTTATGACATTC-aminolinker-3′) and a bridge oligo (for 21 nt small RNA, 5′-GAATGTCATAAGCGACTATACAAGGATCTACCTCA-3′: for 24 nt small RNA, 5′ - GAATGTCATAAGCGGAGACTATACAAGGATCTACCTCT-3′) and annealed in 1× capture buffer (10 mM Tris-HCl (pH 7.5), 75 mM KCl). This was followed by ligation of the guide RNA to the ligation oligo using the DNA Ligation Kit (Takara). Subsequently, unreacted ligation oligo was dephosphorylated using alkaline phosphatase (Calf intestine) (Takara) and formamide dye [10 mM EDTA, pH 8.0, 98% (w/v) deionized formamide, 0.025% (w/v) xylene cyanol and 0.025% (w/v) bromophenol blue] was added. These samples were separated on a 15% denaturing acrylamide gel. Gels were analyzed by PhosphorImager (FLA-7000, Fujifilm Life Sciences). A known concentration of small RNA was subjected to the same reaction, and serial dilutions were performed to create a standard curve.

### 3′ end labeling of target RNA/DNA

Target RNA was labeled at the 3′ end using Poly(A) Polymerase (Takara) and [^32^P]cordycepin-5′-triphosphate. Target DNA was similarly labeled at the 3′ end using Terminal Deoxynucleotidyl Transferase (Affymetrix) and [^32^P]cordycepin-5′-triphosphate. Unreacted cordycepin was then removed using a G25 column (cytiva), followed by purification with phenol-chloroform and chloroform, and the products were analyzed on a 15% denaturing acrylamide gel. The labeled RNA or DNA was excised from the gel and eluted overnight in PK buffer (100 mM Tris-Cl, pH 7.5, 200 mM NaCl, 2 mM EDTA, 1% (w/v) SDS) while rotating. A standard curve was drawn using a dilution series of the labeled RNA/DNA prior to G25 column purification.

### Binding assay

A filter binding assay was conducted following the procedures outlined in previous studies (13, 23). To determine the equilibrium constant, 20 pM of radiolabeled target RNA/DNA was incubated with increasing concentrations of the catalytic-mutant AtAGO-RISC (AtAGO1-, AtAGO2-, AtAGO4-, AtAGO6- and AtAGO9-RISCs) in the equilibration buffer for 1 hour at 25°C. The reaction mixture was then passed through two layers of filters, consisting of a Protran nitrocellulose membrane (Whatman) followed by a Hybond N+ nylon membrane (GE Healthcare Bioscience). Subsequently, the membranes were washed with ice-cold equilibration buffer, air-dried, and the signals were detected using a PhosphorImager (FLA-7000, Fujifilm Life Sciences). Multi Gauge (Fujifilm) was used to quantify the data, and dissociation constants were calculated using Igor64 (WaveMetrics) using the following formula (modified from Wee et al., 2012):

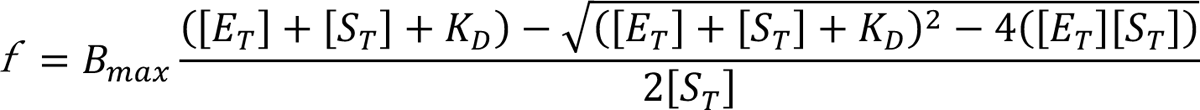

where *f* is the fraction target bound, *B*_max_ is the maximum bound fraction, [*E*_T_] is the total enzyme concentration, [*S*_T_] is the total RNA target concentration, and *K*_D_ is the apparent equilibrium dissociation constant.

### Multiple sequence alignment and protein visualization

Multiple sequence alignment and Tree generation were performed by ClustalW2. Protein structure visualization was performed by PyMol.

## Results

### Clade 1 and 2 RISCs have a higher affinity for RNA than for DNA

AGO1 in clade 1 and AGO2 in clade 2 are known to function via binding to target RNAs in the nucleus as well as in the cytoplasm. However, it remained unclear how RISCs prioritize binding to RNA in the nucleus, where DNA is abundant.

To investigate whether clade 1 and clade 2 RISCs exhibit higher affinity for RNA or DNA binding, I conducted a filter binding assay. Since RISC cleaves the target, making it difficult to accurately determine the equilibrium binding constant, I used catalytic mutant AGOs. AGO proteins were produced in a tobacco cell-free system, and after being programmed with 21-nt small RNAs, they were affinity purified with an N-terminal FLAG tag. Subsequently, RISCs were eluted through SUMOstar protease treatment (23). Quantification of RISCs was performed by quantifying the bound small RNA using the splint ligation method (Supplementary Figure 1). The targets used were 28-nt RNAs and DNAs labeled with ^32^P cordycepin at the 3′ end (Figure 1D). After binding various concentrations of RISCs with 20 pM target nucleic acids, I conducted a dot blot analysis using nitrocellulose and nylon membranes. Targets bound to RISCs were trapped on the nitrocellulose membrane, while unbound targets were retained on the nylon membrane (Figure 1E, Supplementary Figure 2). The ratio (fraction bound) was plotted against RISC concentration to calculate the dissociation constant (Figure 2). AGO1-RISC exhibited a binding affinity of Kd = 12.6 pM for t1U RNA, a target RNA that is fully complementary to positions g2–g21, but bound to t1T, a target DNA with the same complementarity, at Kd = 334.39 pM (Figure 2 and Supplementary Figure 2). Thus, AGO1-RISC exhibited an approximately 25-fold higher affinity for RNA compared to DNA binding. Similarly, the dissociation constants of AGO2-RISC in clade 2 for t1U and t1T were Kd = 7.78 pM and Kd = 4.33 nM (4330 pM) for DNA, respectively (Figure 2 and Supplementary Figure 2), indicating that RNA binds to RISC with an affinity 600 times higher than that of DNA. The target RNA/DNA with a mismatch in the seed region (positions 4- 5) failed to bind to either RISC (Figure 2), indicating that binding between the target nucleic acid and RISC is sequence specific.

**Figure 2.**
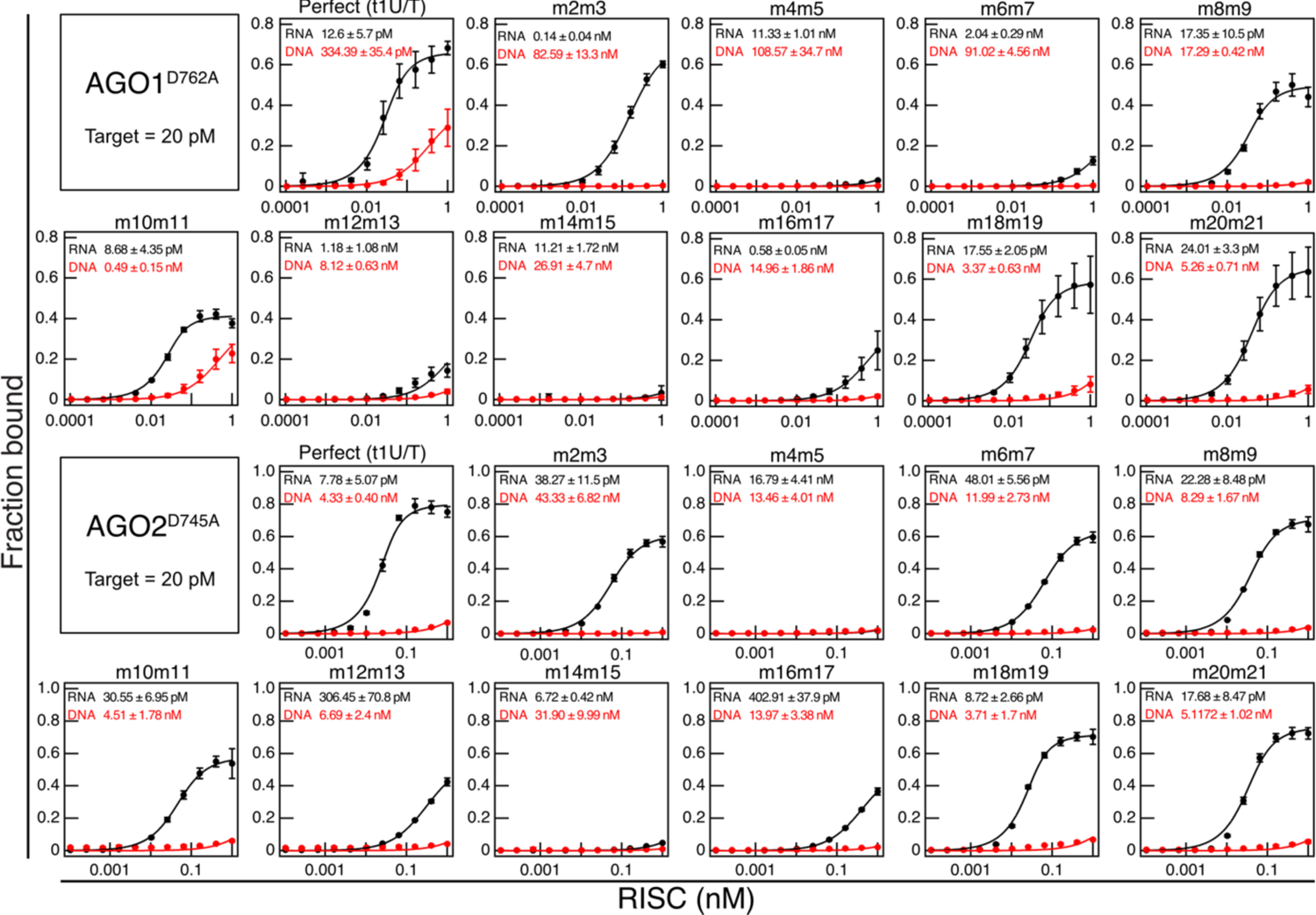
Target binding properties of AGO1- and AGO2-RISCs. Equilibrium binding assay between AGO1/AGO2-RISC and Target RNAs (in black) or DNAs (in red). The sequence of target RNA/DNA are shown in Figure 1D. RISCs were assembled using AGO1^D762A^ and AGO2^D745A^, which harbor mutations in the catalytic active site to prevent cleavage of target nucleic acids. Data represent the mean +/- SD of three independent experiments.

### The 3′ supplementary region is critical for target binding in clade 1 and 2 RISCs

Previously, it was reported that AGO1 and AGO10 require extensive base pairing beyond the seed region for target binding (23, 24, 46). However, the detailed impact of mismatches on target binding remained unclear. Thus, I analyzed the target binding properties of AGO1-RISC and AGO2-RISC using target RNA/DNA with dinucleotide mismatches against the guide strand (Figure 2 and 3). AGO1-RISC exhibited a binding affinity more than 10 times lower when dinucleotide mismatches were introduced in the seed (2–7) region compared to t1U. Particularly, the introduction of mismatches at the 4–5 position resulted in a 1000-fold decrease in affinity compared to t1U. Interestingly, introducing mismatches in the 3′ supplementary region also significantly reduced the binding affinity, and mismatches at the 14–15 position caused a similar 1000-fold reduction in binding affinity as those at the 4–5 position. On the other hand, the introduction of mismatches in the central region and tail region did not affect the binding affinity. In the case of AGO2-RISC target binding, introducing mismatches at positions 4–5 and 14–15 also resulted in a reduction in binding affinity by approximately 1000-fold. Thus, base pairing in the 3′ supplementary region is essential for the stable binding of clade 1 and clade 2 RISCs to target RNA. However, it is important to note that these two RISCs do not exhibit identical target recognition properties, as several discernible differences have been observed. The first distinction lies in the extent of involvement in stable binding due to base pairing at positions 6–7. In the case of AGO1, the introduction of a mismatch at the 6–7 position led to a reduction in binding stability of over 100-fold, while for AGO2, the reduction was only about 7-fold. Furthermore, the introduction of a mismatch at 2–3 positions also had a milder impact on AGO2 compared to AGO1. Taken together, these results suggest that AGO1 requires higher complementarity in target recognition than AGO2.

**Figure 3.**
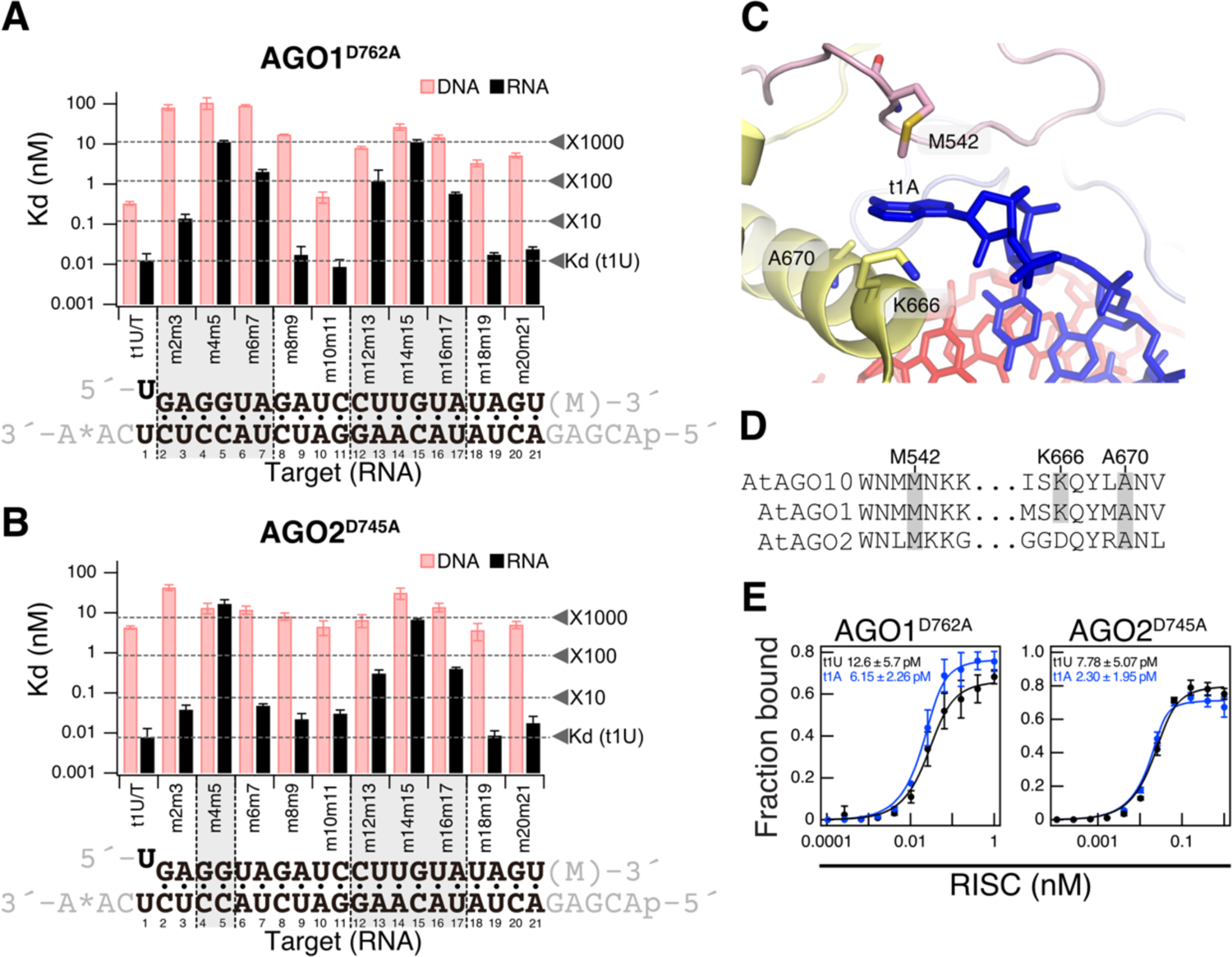
AGO1- and AGO2-RISCs require the 3′ supplementary region in addition to the seed region for target binding. A and B. (top) Dissociation constants (Kd) between AGO1- or AGO2-RISCs and target RNA/DNA with various mutations, calculated from the data in Figure 2. (bottom) Base pairing between guide strand and target RNA (t1U). Gray shading indicates regions where the dissociation constant drops more than tenfold compared to t1U when dinucleotide mismatches are present. C. Structure of AGO10 recognizing t1A [PDB ID: 7SWF]. D. Multiple sequence alignment of AGO10’s amino acid residues involved in t1A interaction (M542, K666, A670) with the corresponding regions in AGO1 and AGO2. E. Equilibrium binding assay using t1A RNA. t1A (Figure 1D) binds to AGO1-RISC and AGO2- RISC with higher affinity than t1U. Data represent the mean +/- SD of three independent experiments. t1U data is the same as in Figure 2.

### The nucleotide at the t1 position of the target RNA affects the binding affinity with RISC

Because the nucleotide at the 5′ end (g1) of the guide strand docks into the pocket of the MID domain, the corresponding nucleotide (t1 nucleotide) in the target strand does not form a base pair (22). However, in animals, AGOs possess a t1-binding pocket, and it is known to enhance the binding between the target and RISC by interacting with adenine (t1A) through water mediation (47). Recently, the structure of *Arabidopsis* AGO10-RISC in complex with the target was elucidated, revealing that residues M542, A670, and K666 are in contact with t1A (Figure 3C) (46). These residues are completely conserved in the same clade 1 AGO1 and are partially conserved in clade 2 AGO2 (Figure 3D). However, the role of the t1 position nucleotide in the interaction between plant RISC and target RNA has not been analyzed to date. Therefore, I investigate the impact of the t1 position nucleotide on the binding between the target and AGO1-RISC and AGO2- RISC by using t1A and t1U RNA. AGO1-RISC bound to t1A with a Kd of 6.15 pM, which was approximately two times higher affinity compared to t1U (Kd = 12.6 pM) (Figure 3E). AGO2- RISC also exhibited approximately three times higher affinity for t1A (Kd = 2.30 pM) compared to t1U (Kd = 7.78 pM) (Figure 3E). These results indicate that the nucleotide at the t1 position influences the interaction with plant RISC, and adenine at the t1 position moderately enhances the affinity between AGO1/AGO2-RISC and target RNA.

### Clade 3 RISC exhibits high affinity binding to DNA

AGO4, AGO6, and AGO9, belonging to clade 3, load 24-nt siRNAs and facilitate DNA methylation. According to the current model, to promote RdDM, clade 3 AGOs are believed to bind to the nascent transcript produced by the plant-specific RNA polymerase Pol V (26). Indeed, it has been observed that transcripts generated by Pol V are cleaved by AGO4 (48), providing further support for the Pol V RNA-based RdDM model. However, there have been a report suggesting the possibility of AGO4-RISC acting by binding to DNA (49). Yet, it remained unclear whether this binding occurs through base pairing or involves non-base pairing interactions with nearby DNA. Therefore, I prepared purified AGO4-RISC, AGO6-RISC, and AGO9-RISC using the same method as AGO1 and AGO2-RISC and analyzed the affinity between target RNA/DNA and clade 3 RISCs through filter binding assays. For guide RNAs, I used 24-nt small RNAs. As expected, AGO4-RISC, AGO6-RISC, and AGO9-RISC showed high affinity for RNA (AGO4- RISC: t1U, Kd = 1.71 pM; AGO6-RISC: t1U, Kd = 50.36 pM; AGO9-RISC: t1U, Kd = 12.72 pM) (Figure 4 and Supplementary Figure 2). Interestingly, they exhibited equivalent or even higher affinity for DNA (AGO4-RISC: t1T, Kd = 0.93 pM; AGO6-RISC: t1T, Kd = 7.30 pM; AGO9-RISC: t1T, Kd = 2.12 pM) (Figure 4 and Supplementary Figure 2). It is noteworthy that these bindings are sequence-specific. Target RNA/DNA with mismatches introduced at the seed region positions 4–5 barely bound to any RISC (Figure 4). Thus, clade 3 RISC, unlike clade 1 and 2 RISC, does not avoid binding to DNA and, in fact, prefers it over RNA.

**Figure 4.**
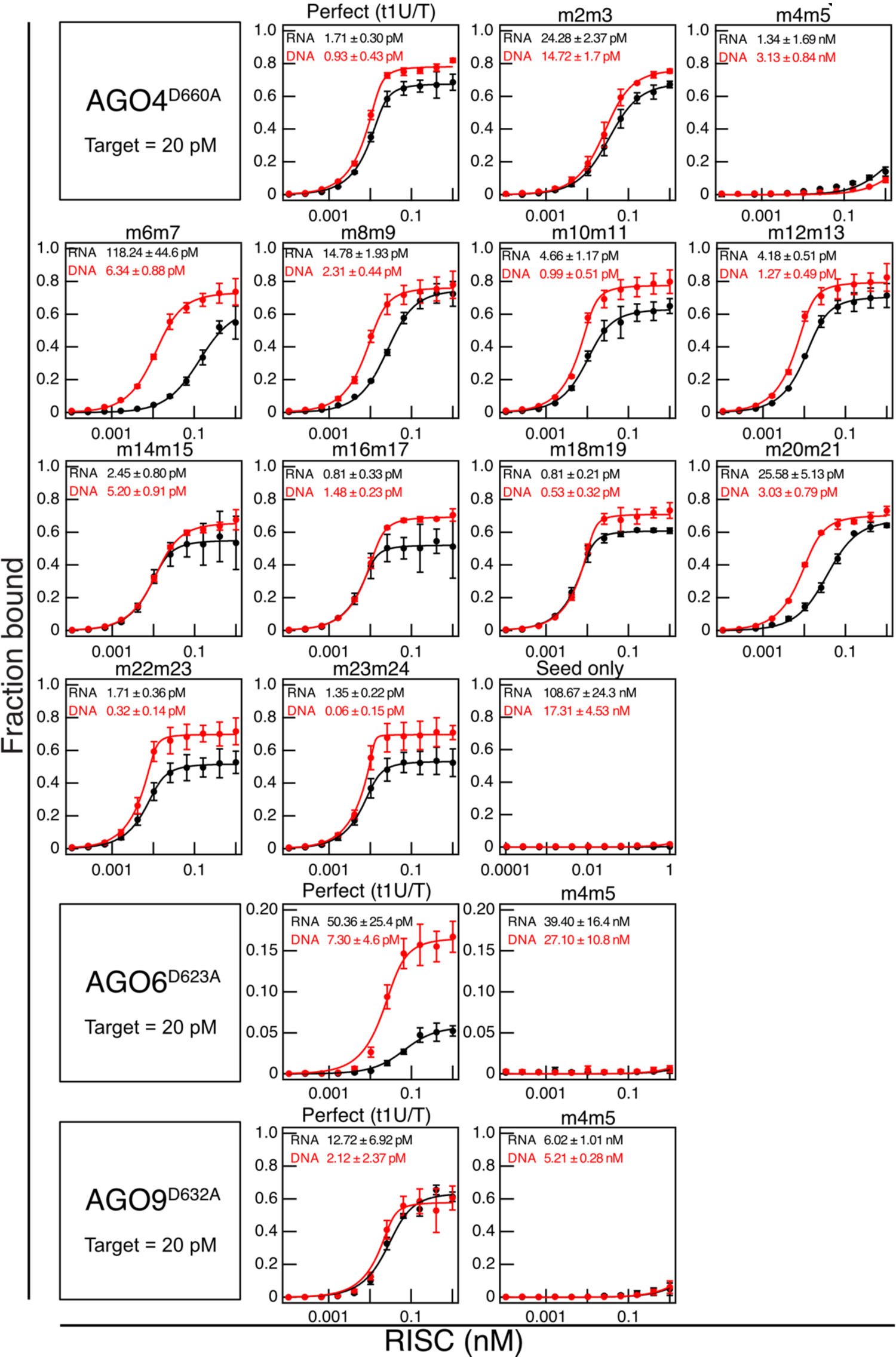
Clade 3-RISCs bind to DNA with an affinity equal to or greater than that of RNA. Equilibrium binding assay between clade 3-RISCs and target RNAs (in black) or DNAs (in red). The sequence of target RNA/DNA are shown in Figure 1D. RISCs were formed using AGO4^D660A^, AGO6^D623A^, and AGO9^D632A^, which harbor mutations in the catalytic active site to prevent cleavage of target nucleic acids. Data represent the mean +/- SD of three independent experiments.

### AGO4-RISC tolerates mismatches beyond the seed region in target binding

Until now, the target recognition mechanism of clade 3 RISC was largely unknown, leaving the extent to which the RdDM mechanism can act in a trans manner unclear. Therefore, I conducted an analysis of the target recognition properties of AGO4-RISC using target RNA/DNA with dinucleotide mismatches against the 24-nt guide strand (Figure 1D, 4 and 5). Similar to AGO1- RISC and AGO2-RISC, AGO4-RISC showed a significant decrease in binding affinity when mismatches were present in the seed region, especially at positions 4–5. However, unlike AGO1- and AGO2-RISCs, the 3′ supplementary region is not essential for target binding in AGO4-RISC, and the only region that substantially reduced the binding affinity, by approximately 10-fold, was limited to positions 20–21, apart from 4–5 (Figure 4 and 5). In other words, AGO4-RISC demonstrated stable binding to the target even when mismatches were present beyond the seed region. Importantly, even in the presence of mismatches, AGO4-RISC generally exhibited stronger binding to DNA than RNA. Particularly, when mismatches were present at positions 6–7, 8–9, 20– 21, and 23–24, the binding to DNA was approximately 7 to 20 times stronger than that to RNA (Figure 4 and 5).

**Figure 5.**
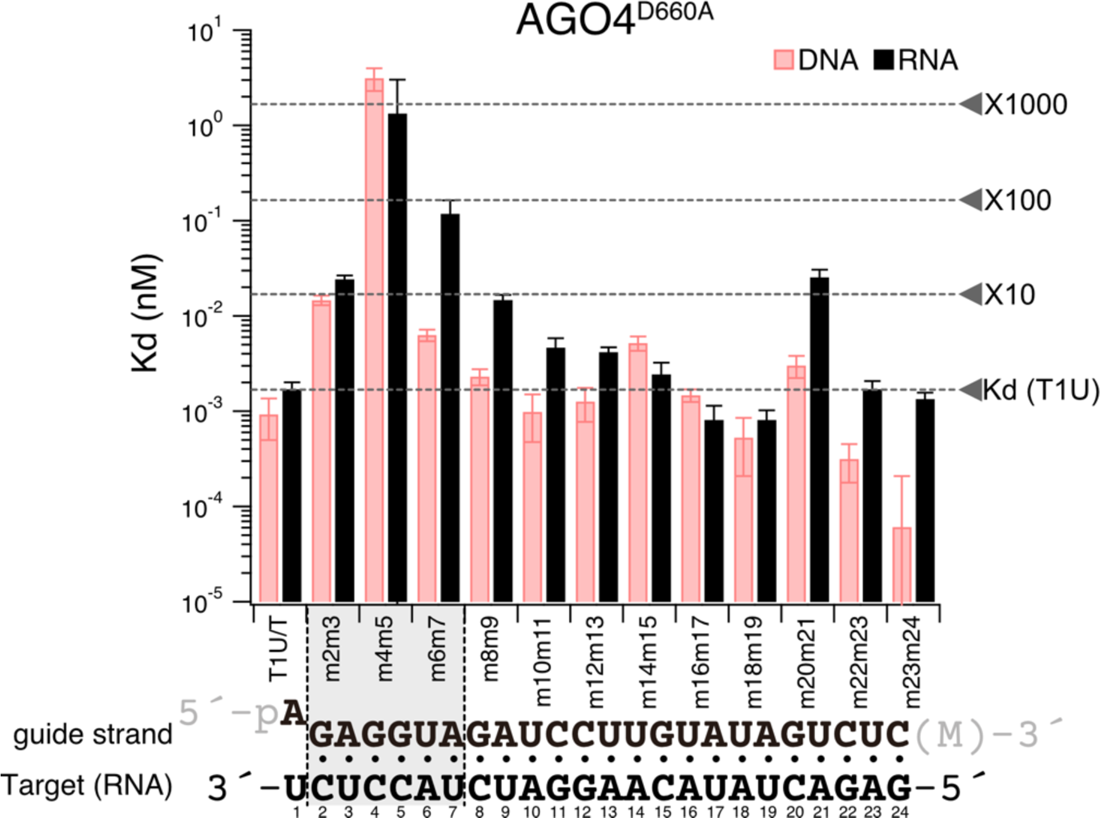
AGO4-RISC tolerates mismatches beyond the seed region in target binding. (top) Dissociation constants between AGO4-RISC and target RNA/DNA with various mutations, calculated from the data in Figure 4. (bottom) Base pairing between guide strand and target RNA (t1U). Gray shading indicates regions where the dissociation constant drops more than tenfold compared to t1U when dinucleotide mismatches are present.

Binding affinity showed only a mild decrease despite mismatches beyond the seed region. From this observation, I hypothesized that AGO4-RISC, like metazoan miRNA-RISC, may stably bind to its target through base pairing only in the seed region. I conducted a filter binding assay using target RNA/DNA with complementarity only in positions 2–8 (Figure 1D). Contrary to my expectations, AGO4-RISC did not bind to this target at all. In other words, while AGO4-RISC can tolerate dinucleotide mismatches beyond the seed region, it is unstable with the seed alone, emphasizing the necessity of additional base pairing for stable binding to the target.

### Cytoplasmic RISC differentiates between RNA and DNA at the 3′ supplementary region

In this study, I have revealed that clade 3 nuclear RISCs bind to DNA with an affinity equivalent to or even higher than that of RNA. On the other hand, clade 1 and clade 2 cytoplasmic RISCs avoid DNA and exhibit high affinity binding to RNA. Now, the question arises: How do cytoplasmic RISCs distinguish between DNA and RNA? To address this question, I performed *in vitro* binding assays using two chimeric targets with mixed DNA and RNA sequences. The target nucleic acids t2–t8_RNA and t9–t24_RNA present distinct structural features: t2–t8_RNA encompasses RNA in the seed region (positions t2 to t8) and DNA elsewhere, whereas t9– t24_RNA incorporates RNA beyond the seed region (positions t9 to t24), with the seed region and remaining sequence being DNA. The AGO4-RISC bound both targets with high affinity (t2– t8_RNA, Kd = 0.87 pM; t9–t24_RNA, Kd = 0.75 pM). However, the AGO1-RISC displayed a contrasting binding profile, showing high affinity for t9–t24_RNA (Kd = 0.28 pM) but a roughly 400-fold lower affinity for t2–t8 RNA (Kd = 118 pM). This suggests that the cytoplasmic RISC can maintain high-affinity interactions with targets even if the seed region is composed of DNA, but its affinity drops when regions beyond the seed are DNA. Given that neither central nor tail regions affect target binding, I deduce that the cytoplasmic RISC differentiates between RNA and DNA at the 3′ supplementary region, avoiding DNA interactions.

## Discussion

### The necessity of base pairing beyond the seed region in target binding of plant RISCs

Metazoan miRNA-RISCs (e.g., HsAgo1–4 and DmAgo1) are known to rely on base pairing in the seed region alone for target binding. However, it has been demonstrated that clade 1 AGO1- and AGO10-RISCs necessitate base pairing beyond the seed region (23, 24). In this study, it was uncovered that not only clade 1 RISCs but also a clade 2 RISC demand base pairing in the 3′ supplementary region, particularly at positions 14–15, for target binding (Figure 2 and 3). Additionally, it has become evident that clade 3 AGO4-RISC cannot bind to the target with seed pairing alone (Figure 4). These results suggest that plant AGOs, unlike metazoan AGOs, weaken seed pairing formation. A previous structural study of AGO10-RISC suggested that 1) the PIWI domain’s loop (PIWI-loop) of AGO10 contacts the L2-loop, inhibiting 3′-side seed pairing (position 6–7), thus weakening seed pairing; and 2) extended base pairing beyond the seed region induces a conformational change in AGO10, disrupting the physical contact between the PIWI-loop and L2-loop, which, in turn, promotes complete base pairing in the seed region (24) (Figure 6C: RNA target). Given the high sequence identity of AGO1’s PIWI loop with that of AGO10, it is possible that AGO1 operates with a similar mechanism in target recognition. On the other hand, the PIWI-loops of AGO2 and AGO4 exhibit lower sequence similarity with AtAGO10’s PIWI-loop (Supplementary Figure 3), suggesting that the seed weakening mechanism in clade 2 and clade 3 RISCs may not be the same as those in clade 1-RISCs. To elucidate these mechanisms, structural and biochemical analyses of target-bound RISCs in clade 2 and 3 will be needed.

**Figure 6.**
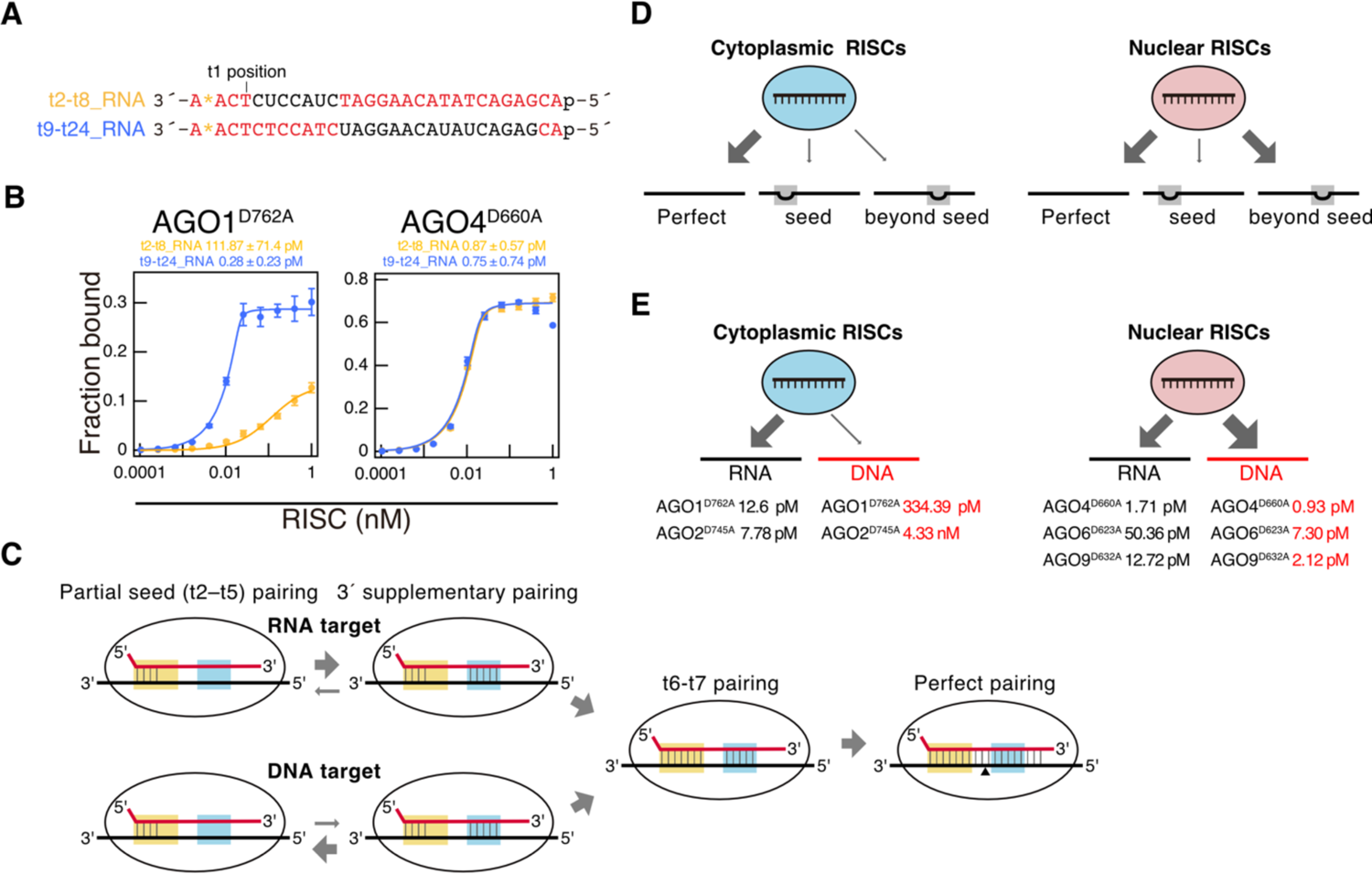
Cytoplasmic RISC distinguishes between RNA and DNA in the 3′ supplementary region, avoiding binding to DNA. A. Sequences of RNA/DNA chimeric targets. t2–t8_RNA is a chimeric nucleic acid in which positions t2 to t8 are RNA and the rest is DNA. t9–t24_RNA is a chimeric nucleic acid in which positions t9 to t24 are RNA and the rest is DNA. A. Equilibrium binding assay with RNA/DNA chimeric targets. AGO4-RISC shows equivalent binding affinity to both targets, while AGO1-RISC binds with high affinity only to t2–t8_RNA. The binding assay between t9–t24 RNA and AGO4-RISC showed a decrease in fraction bound when the RISC concentration exceeded 0.4 nM. Therefore, values below 0.4 nM RISC concentration were utilized for fitting in this study. A. Model for the mechanism by which cytoplasmic RISCs avoid binding to DNA and bind with high affinity to RNA. Initially, the 5′ side of the seed region (g2–g5) binds weakly with the target RNA/DNA, without discrimination between RNA and DNA. Next, if the target is RNA, base pairing in the 3′ supplementary region promotes base pairing at the 3′ side of the seed region (position 6–7), resulting in the completion of full seed pairing. Conversely, if the target is DNA, the shape of the duplex differs from the A-form and clashes with the AGO protein, thus preventing base pairing in the 3′ supplementary region. As a result, base pairing at the 3′ side of the seed region (position 6–7) does not form, leading to very weak affinity. A. Cytoplasmic RISCs require extensive base pairing in the seed and beyond seed regions for stable binding, whereas nuclear RISCs tolerate dinucleotide mismatches in the beyond seed region. A. Cytoplasmic RISCs avoid binding to DNA and preferentially bind to RNA, whereas nuclear RISCs bind to DNA with equal or higher affinity than to RNA.

### Physiological role of mismatch tolerance in target binding of plant RISCs

AGO4-RISC can tolerate dinucleotide mismatches outside the seed region for target binding (Figure 6D). The wide range of mismatch tolerance in this region may have a role in compensating for the error-prone transcription of RNA polymerase IV (pol IV), which generates the precursor of the 24-nt siRNA (50). In addition, this mismatch tolerance demonstrates the potential of clade 3 RISC to induce RdDM not only in *cis* but also in *trans*. Indeed, it has been reported that small RNAs with imperfect complementarity to their targets can trigger stable and heritable RdDM (51). Transposable elements (TEs) are prone to accumulating mutations, but this property of mismatch tolerance may enable even 24-nt siRNAs derived from highly mutated ancient TEs to effectively inhibit active TEs. Interestingly, this mechanism of tolerating mismatches beyond the seed region is also utilized by the piRNA-PIWI-RISC pathway in the germline of animals (52, 53). While the piRNA pathway and the RdDM pathway have evolved independently, they appear to have converged in allowing and functioning with mismatches to efficiently suppress highly mutated TEs and viruses.

In comparison to AGO1, which incorporates miRNAs for stringent gene expression regulation, AGO2, functioning in antiviral responses, exhibits tolerance to dinucleotide mismatches at positions t2–t3 and t6–t7 (Figure 2 and 3). This suggests a potential adaptation in the target recognition mechanism of AGO2, possibly to accommodate the high frequency of mutations associated with RNA viruses.

### Distinguishing mechanisms between DNA and RNA targets in cytoplasmic RISC and its biological implications

The experiments involving RNA/DNA chimera targets in this study suggest that cytoplasmic RISC distinguishes between DNA and RNA targets in the 3′ supplementary region (Figure 6). Now, how do cytoplasmic RISCs differentiate between DNA and RNA in the 3′ supplementary region? The structure of AGO10 showed that a loop in the PIWI domain, referred to as conserved sequence 7 (cS7), interacts with the minor groove of the guide RNA-target RNA duplex at positions 12–14. Additionally, the backbone of positions t13 and t14 is contacted by the L1 hairpin (46). This indicates that clade 1 AGO10 rigorously licenses RNA-RNA duplexes. On the other hand, RNA-DNA heteroduplexes, while resembling RNA-RNA A-form duplexes, possess a minor groove approximately ∼2Å narrower (54). Furthermore, the ternary complexes of prokaryotic AGO-small RNA-target DNA exhibit variations in helix positioning, pitch, and groove depth, presenting a structure entirely distinct from the A-form of RNA duplexes (55, 56). Combining these insights, a model emerges where RISCs of clade 1 and 2 can bind to target DNA with partial seed pairing; however, elements like cS7 and the L1 hairpin could impede stable DNA-RNA duplex formation in the 3′ supplementary region. This could make the completion of seed pairing challenging, leading to unstable target binding (Figure 6C: DNA targeting).

Several biological rationales can be posited for why cytoplasmic AGO proteins distinguish between RNA and DNA, showing a preference for RNA binding. It is known that AGO1 shuttles between the nucleus and the cytoplasm, and there are reports suggesting that the formation of RISC with miRNAs takes place within the nucleus (57). If AGO1 were to bind DNA with high affinity, it could inadvertently interact with miRNA genes, potentially inhibiting the transcription of miRNA precursors and consequently decreasing the availability of mature miRNAs. Alternatively, DNA binding could result in the retention of mature RISCs within the nucleus, thereby diminishing the concentration of functional RISCs in the cytoplasm. To avert these outcomes, it is possible that AGO proteins have evolved to maintain low DNA binding affinity while preferentially binding to RNA. On the other hand, the evolution of clade 1 and 2 AGOs may not have been driven by an active avoidance of DNA interactions. Rather, these proteins could have evolved to bind more strongly to RNA, resulting in an inability to accommodate the structurally distinct RNA-DNA heteroduplexes within the RISC.

### The mechanism and potential roles of clade 3 RISCs in accepting DNA targets

All clade 3 RISCs exhibited target DNA binding affinities equivalent to or greater than those for target RNA. Remarkably, in the presence of mismatches at positions 6–7 and 23–24, AGO4-RISC exhibited an approximately 20-fold higher affinity for DNA targets than for RNA (Figure 4). As previously noted, the DNA-RNA heteroduplex has a different shape compared to the RNA’s A-form duplex, necessitating clade 3 AGO to maintain a unique structural conformation to bind with high affinity to DNA. In AGO4, 6, and 9 from clade 3, a characteristic feature is observed where glycine, an α-helix breaker within the cS7, is substituted by serine. Moreover, in the L1 loop, a glycine and arginine pair, which is highly conserved in many AGO and PIWI clade proteins and makes contact with the backbones of t13 and t14, is altered in AGO4, 6, and 9, with each being replaced by a pair of arginine and glutamine, respectively (Supplementary Figure 4). It is possible that these differences in the loops play a role in accommodating DNA-RNA heteroduplexes.

Clade 3 RISC is believed to promote RdDM by binding to Pol V’s C-terminal domain and nascent transcript by recruiting the *de novo* DNA methyltransferase DRM2. The interaction between AGO4 and Pol V’s transcript is evident from the AGO4-dependent cleavage of Pol V transcripts (48), supporting the Pol V RNA-guided RdDM model. However, it has not been ruled out that AGO4-RISC may also bind to single-stranded DNA regions that could potentially arise during Pol V transcription and facilitate RdDM. In previous studies, the use of a UV laser-based crosslink immunoprecipitation assay demonstrated the direct binding of AGO4 to target DNA regions (49). While these results alone could not rule out the possibility of base-pairing independent binding of AGO4 to target DNA, the addition of data from the current study, showing strong binding of all three clade 3 RISCs to DNA, may warrant serious consideration of a model in which RdDM is facilitated through base-pairing interactions between RISC and target DNA in addition to those between RISC and target RNA. Furthermore, it is known that the Arabidopsis genome frequently forms R-loops (58). Given the strong binding affinity between clade 3 RISC and DNA, it is quite possible that these RISCs bind directly to exposed single-stranded DNA regions to perform functions beyond RdDM. Moreover, the known interactions between prokaryotic AGOs and phage DNA (2) open up the possibility that eukaryotic nuclear-RISCs, through direct DNA binding, could play a crucial role in hindering the proliferation of DNA viruses. Therefore, delving into the biological significance of the high DNA binding affinity of clade 3 RISCs has the potential to reveal new mechanisms of small RNA-mediated regulation of gene expression in eukaryotes.

## Data availability

The data underlying this article are available in the article and in its online supplementary material.

## Author contributions

Hiro-oki Iwakawa: Conceptualization, Formal Analysis, Investigation, Visualization, Writing – original draft.

## Acknowledgements

I thank Yuichiro Watanabe and Atsushi Takeda for providing plasmids encoding AtAGO4, AtAGO6, and AtAGO9, Wei Liu for providing pBYL-3×FLAG-SUMO-AGO6^D623A^ and pBYL- 3×FLAG-SUMO-AGO9^D632A^, and Kaori Kiyokawa for experimental assistance. I also thank Dr. Yukihide Tomari, Dr. Masahiro Naganuma and members of the Iwakawa laboratory for discussions and critical reading of the manuscript.

