## Supplementary information for "The clade-specific target recognition mechanisms of plant RISCs"

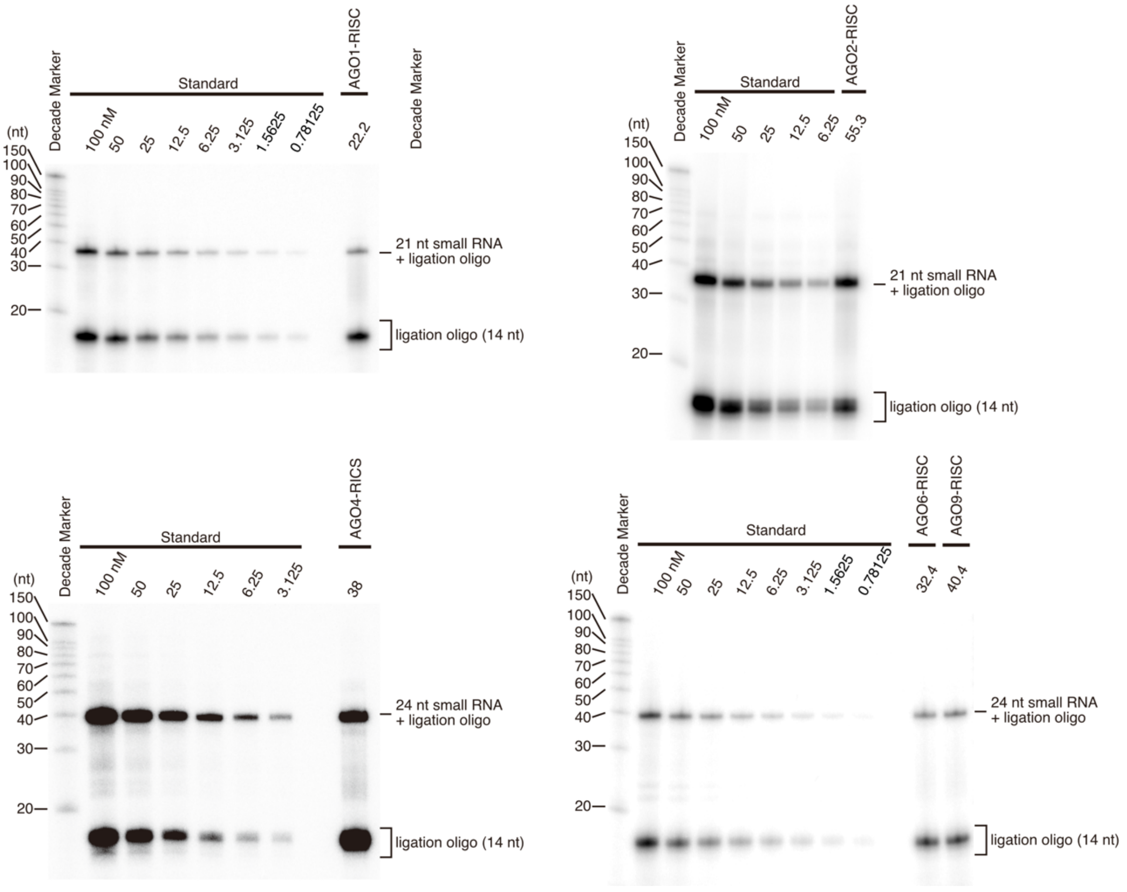


**Supplementary Figure 1. Quantification of RISC using the splint ligation method.**

After deproteinization of the purified RISC, the guide strand and a radiolabeled ligation oligo were brought into proximity using a bridge oligo, followed by ligation (21/24-nt small RNA + ligation oligo). Unreacted ligation oligo, which was not completely dephosphorylated by alkaline phosphatase, is observed at the lower part of the gel. A known concentration of small RNA was subjected to the same reaction, and serial dilutions were performed to create a standard curve.

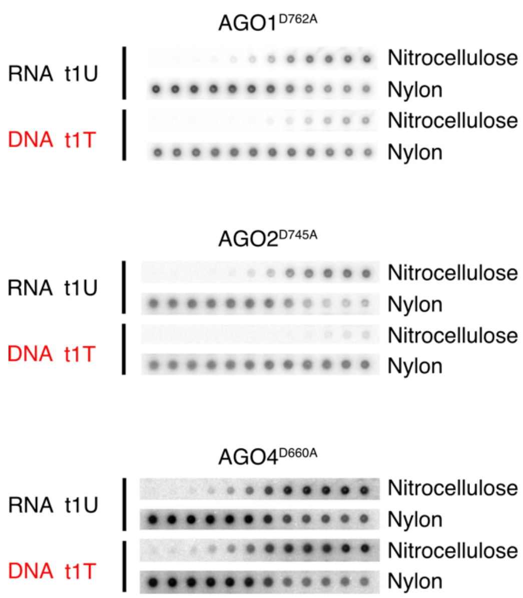


**Supplementary Figure 2. Equilibrium binding assay of t1U/T to RISC.**

RISC-bound target is blotted onto the nitrocellulose membrane and unbound target RNA is blotted onto the nylon membrane.

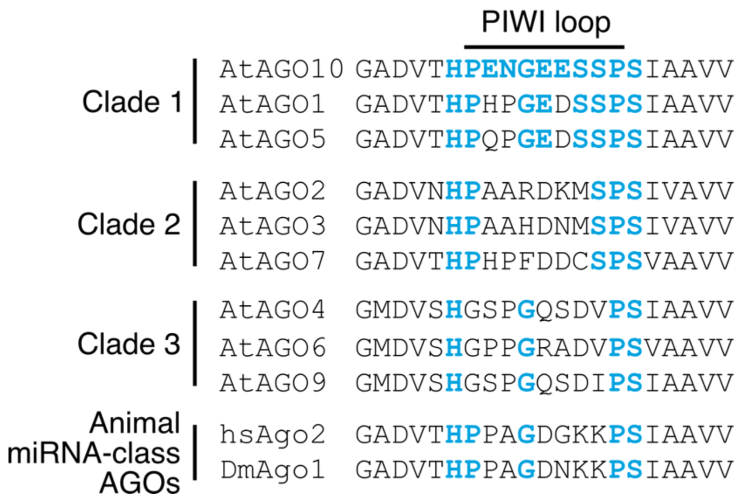


**Supplementary Figure 3. Multiple alignment of amino acids in the PIWI loop region in *Arabidopsis* AGOs and animal AGOs.**

Bold blue letters indicate amino acids identical to the PIWI loop in AGO10.

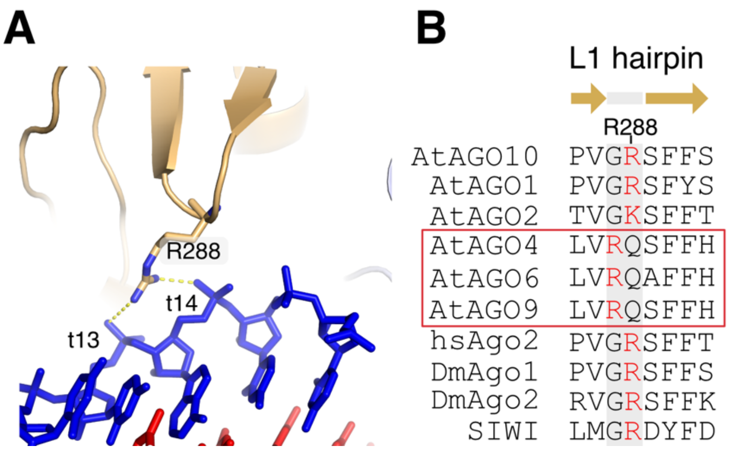


**Supplementary Figure 4. The L1 hairpin contacts the target RNA in the 3′ supplementary region.**

A. The structure of L1 hairpin in the AGO10-miRNA-target complex [PDB ID: 7SWF]. The arginine residue in the L1 hairpin contacts the backbone of t13 and t14 of the target RNA.

B. Multiple alignments of the amino acid sequence of the L1 hairpin. The turn of the hairpin is formed by different amino acids only in clade 3 AGOs.

```
                            21U-guide
                     pUGAGGUAGAUCCUUGUAUAGU(M)
21U duplex              ||| |||| |||||||||
                 (M)UCUCUCGAUCUUGGAACAUAUp

                            21A-guide
                     pAGAGGUAGAUCCUUGUAUAGU(M)
21A duplex              ||| |||| |||||||||
                 (M)UCCCUCGAUCUUGGAACAUAUp

                            24A-guide
                     pAGAGGUAGAUCCUUGUAUAGUCUC(M)
24A duplex             |||| ||| |||||| ||||||
                 (M)UCUCUCUAUCAAGGAACUUAUCAGp
```

**Supplementary Figure 5. Small RNA duplexes used in this study.**

The 3′ end of small RNA is modified with 2′-O-methyl (M).
